# Closing the Sim-to-Real Gap: An End-to-End Robotic Ultrasound System Leveraging In Vivo Reinforcement Learning and 3D-Prior Guided Hybrid Control

**DOI:** 10.64898/2025.12.17.694846

**Authors:** Hui Tang, Chenxi Xie, Boyang Zhou, Yikang Sun, Qi Yan, Dun Xue, Jieyuan Hu, Fangzi Liu, Qinglin Liu, Xinyuan Hu, Li Wu, Yi Wang, Huixiong Xu, Meng Yang

## Abstract

Abdominal ultrasound is a crucial first-line diagnostic tool, yet its efficacy is inherently constrained by a strong dependency on operator skill, leading to significant inter-operator variability and limiting its widespread adoption. While robotic ultrasound systems with artificial intelligence have emerged to mitigate this, current approaches face critical limitations: they often rely on simulated or phantom data for training, which hampers generalizability, and employ control strategies that lack the flexibility to adapt to complex human anatomy. To address these challenges, this paper introduces a novel, end-to-end robotic system for autonomous abdominal scanning. Our methodology is systematic: it begins with coarse scan planning via RGB and 3D point cloud-based localization of key anatomical landmarks (e.g., xiphoid process). The core of our approach involves training a reinforcement learning scanning policy directly on live human volunteers, enabling the development of a strategy that is robust to anatomical diversity. This high-level policy is executed by a sophisticated hybrid force-position controller, enhanced with real-time normal vector calibration and 3D point cloud-based respiratory phase detection to dynamically adjust contact force, particularly during inspiration. This innovation proves critical for improving scan quality in subjects with higher Body Mass Index (BMI). Extensive validation against expert sonographers demonstrates that our system achieves high precision in standard plane recognition and superior scanning efficiency. Furthermore, to holistically assess performance where standard planes are difficult to acquire, we introduce a 3D reconstruction-based coverage metric. Results confirm that our system delivers strong adaptability and significantly improved consistency, marking a substantial step toward clinically viable, operator-independent ultrasound robotics.

## Introduction

Medical imaging is indispensable in modern clinical practice, with ultrasound (US) standing out as a vital, first-line diagnostic tool due to its real-time, radiation-free, and portable nature. It has become an essential and often preferred modality for real-time examination, particularly for abdominal and emergency assessments [6]. However, the widespread adoption of ultrasound is constrained by significant operational challenges. Unlike static imaging technologies such as CT and MRI, diagnostic ultrasound is a dynamic process that critically depends on the sonographer’s skilled coordination of “hand-eye-brain” [1]. This strong reliance on operator expertise inevitably introduces subjective variability and inconsistency in both image acquisition quality and diagnostic interpretation [1][4][5]. Compounding this issue is the global shortage of healthcare professionals proficient in ultrasonography. This scarcity, coupled with inherent inconsistencies in manual scanning, creates a pressing clinical dilemma: an urgent need to augment or even substitute human operators to ensure standardized, accessible, and high-quality ultrasound diagnostics [3].

A promising direction involves the use of artificial intelligence to guide the probe. For instance, some systems employ an artificial neural network that integrates real-time ultrasound video with motion data from a probe-mounted inertial measurement unit (IMU) to predict optimal guidance signals, thereby demonstrably lowering the required level of operator expertise [3]. This approach aligns with the encouraging concept of “Abdominal AI,” which refers to the application of artificial intelligence to assist in acquiring, interpreting, and standardizing ultrasound examinations of the abdominal region [6]. By facilitating the consistent capture of key anatomical planes—such as the longitudinal kidney, transverse gallbladder, and subcostal liver views—this technology directly tackles the challenge of scanning consistency for complex abdominal structures [6].

The pursuit of intelligent, adaptive probe navigation has spurred notable innovation, particularly through the adoption of advanced machine learning paradigms. Encouragingly, reinforcement learning (RL) and imitation learning are gaining traction for teaching robots to perform ultrasound scanning. Frameworks include simulation-based RL trained on binary vessel images that can transfer to real-world scenarios [4], Deep Q-Networks (DQN) for probe position optimization [5], and methods like inverse reinforcement learning (IRL) that learn scanning policies from expert demonstrations [8][14][15]. These pioneering works often discretize probe movements into a set of actions, showcasing a promising trend toward data-driven, autonomous scanning strategies.

Despite these encouraging advances, a significant critical flaw persists in the current research landscape: a heavy reliance on simulated and phantom data for training and validation. Many of these sophisticated learning models, including DQN [5] and imitation learning systems [8], are trained primarily on datasets collected from phantoms or simulations [9][10][11]. While this is a necessary preliminary step, it severely limits the models’ generalizability and robustness when confronted with the vast anatomical diversity and physiological complexity of live human patients.

Further complicating the development of robust robotic ultrasound is a notable isolation in control strategy design. While the necessity of both force control (to ensure safe and effective skin contact) and position control (for accurate probe navigation) is universally acknowledged, their seamless integration remains a challenge. Many systems implement only one aspect—for example, employing impedance control for a single degree of freedom [5][7] or relying on position control alone [8]. Although a few systems have implemented force-position hybrid control, their application is often limited to rigid structures like bones and lacks flexibility. These systems typically depend on complex pre-operative path planning and surface mesh reconstruction, requiring a user to pre-select a Region of Interest (ROI) for a predefined scanning path [9][10][12]. This approach is critically lacking in flexibility and does not synergize well with the adaptive, learning-based navigation methods discussed previously, creating a disjointed system architecture.

Amid these challenges, several technologies offer promising pathways for convergence and improvement. A notable and encouraging development is the integration of 3D reconstruction and point cloud technologies for probe navigation. For instance, some systems perform surface reconstruction to handle 3D data and soft tissue displacement for applications like liver tumor tracking [7], while others utilize point clouds to define ROIs and integrate them with motion control algorithms [9]. Although these methods sometimes still rely on predefined positions, their ability to model complex patient-specific anatomy in three dimensions presents a significant opportunity. This technological foundation holds great potential for providing the rich, spatial context necessary to bridge the gap between data-driven navigation and physical, contact-aware control.

Based on the comprehensive analysis of existing limitations, this paper proposes a systematic and effective solution designed to overcome the aforementioned challenges. Our integrated framework progresses from initial landmark localization and coarse scan path definition, to training a reinforcement learning (RL) scanning policy in vivo on real volunteers, down to a lower-level hybrid force-position controller, ultimately converging on the acquisition of high-quality scan data and 3D reconstruction of standard planes. This enables deep, comprehensive scanning of abdominal organs, with a primary focus on the liver. Comparative evaluations with experienced sonographers on scanning efficiency and standard plane recognition accuracy demonstrate that our system exhibits strong adaptability to different subjects and achieves high precision in identifying standard anatomical views. Recognizing that standard plane acquisition rate is an incomplete metric—especially for subjects with higher BMI where certain views are inherently difficult to obtain—our method introduces a complementary evaluation standard. We perform a comprehensive 3D reconstruction of all scanned tissues to calculate the scan coverage ratio. This quantitative measure of anatomical structure visualization breadth serves as a robust and fair supplementary metric for assessing scanning thoroughness, ensuring a holistic performance evaluation beyond the binary success/failure of capturing specific standard planes. This end-to-end pipeline, from intelligent initialization to multi-faceted evaluation, represents a significant stride toward a truly adaptive, clinically viable, and systematically validated robotic ultrasound system for abdominal examination.

## Results

### Overview

The system initiation involves the precise localization of key anatomical landmarks—such as the xiphoid process and umbilicus—in conjunction with the identification of optimal scan starting points. This is accomplished by fusing 3D point cloud data with an RGB-based model, providing a robust spatial context for the subsequent scanning sequence.

A critically novel aspect of our methodology is the training of the RL scanning policy directly on real human volunteers, rather than in simulation or on phantoms. This approach allows the agent to learn a highly adaptive scanning strategy that generalizes across the inherent anatomical variations present in a live population, directly addressing the primary shortfall of phantom-dependent training identified in prior work [5][8][9][10][11].

The learned high-level navigation policy is seamlessly executed by a sophisticated hybrid force-position controller. This controller is enhanced with two key innovations: real-time normal vector calibration along the scan path to maintain optimal probe contact, and the integration of inspiratory phase detection based on 3D point cloud dynamics, which triggers controlled force augmentation. Our experiments substantiate that this integrated control strategy provides a substantial improvement in the successful acquisition rate of standard planes, particularly for volunteers with a higher Body Mass Index (BMI).

### Xiphoid process and the navel detection

A key point detection model was employed to identify the coordinates of the xiphoid process and the navel in RGB images. Subsequently, depth camera data were utilized to obtain point cloud information, allowing the conversion of these two points from two-dimensional image coordinates into three-dimensional coordinates in the working coordinate system.

### Determination of the scanning region and starting points

The scanning region was defined using the coordinates of the xiphoid process and the umbilicus, together with the body contour identified from point cloud data. A costal margin line was constructed by connecting the xiphoid process to the point on the body contour whose y-axis coordinate matched that of the umbilicus. This costal margin line was then translated downward by 6 cm to obtain a parallel reference line. Next, a vertical line was drawn through the xiphoid process and perpendicular to the costal margin line. The quadrilateral formed by these three lines and the body contour was designated as the scanning region.

For probe placement, the starting points for the scanning of standard planes 1, 2, 3, and 7 were set at the xiphoid process. The starting points for standard planes 5 and 6 were defined at the location one-fifth of the total length of the costal margin line segment, measured from the xiphoid process. The starting point for standard plane 8 was defined at the point four-fifths along the same line segment.

### Scanning of the left hepatic lobe and intercostal regions

To ensure comprehensive coverage of the entire liver, we incorporated additional scanning of the left hepatic lobe and intercostal regions into the protocol. The overall scanning procedure begins at the xiphoid process and proceeds downward along the costal margin, sequentially acquiring standard planes 1 through 6. Following completion of standard plane 6, the probe continues with plane 7, then advances to the intercostal scanning sequence performed from superior to inferior, and finally acquires plane 8 (hepatorenal view) before termination. The eight standard hepatic ultrasound planes targeted by the system, along with their clinical significance, are detailed in Table 1.

**Table 1.**
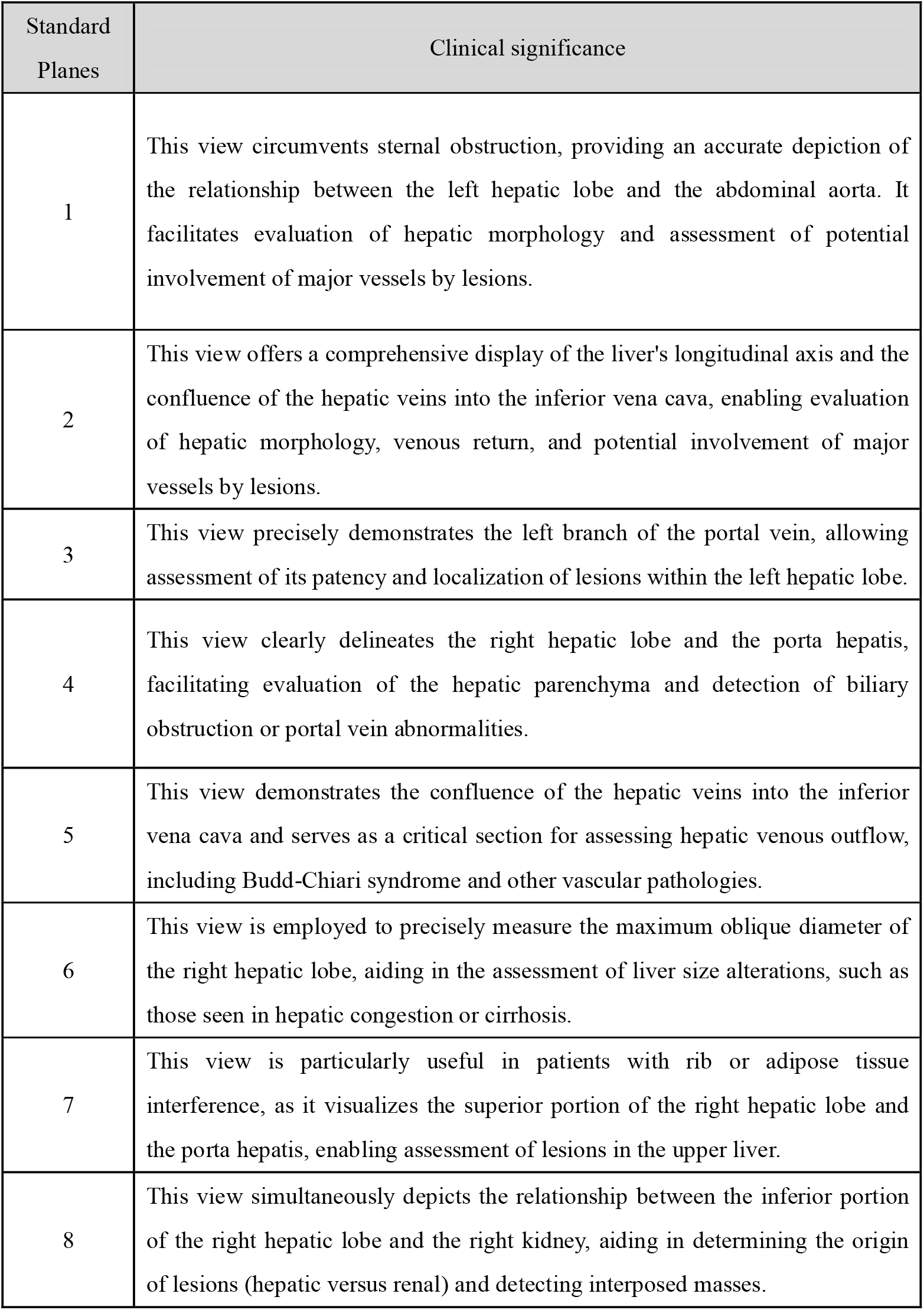
Standard hepatic ultrasound planes targeted by the robotic system and their clinical significance.

Left hepatic lobe scanning was introduced immediately after completion of the scanning of standard plane 1. Especially, seven predefined probe poses were established, starting from the terminal position of standard plane 1. At each pose, only the probe’s Rx angle was decreased by 6°, while all other coordinates remained fixed. This step was repeated seven times, enabling the probe to reach the series of seven positions and thereby complete scanning of the left hepatic lobe.

### Ablation tests

We train the reinforcement learning scanning policy actor using the typical Soft Actor-Critic framework [18]. We trained the model from scratch, without any pre-training that would bias the policy network with expert data. This approach is analogous to training a Go master without human knowledge [19]. With no phantom data or readily collected expert ultrasound image data, we directly train on volunteers from scratch (Fig. 2 b).

**Figure 1:**
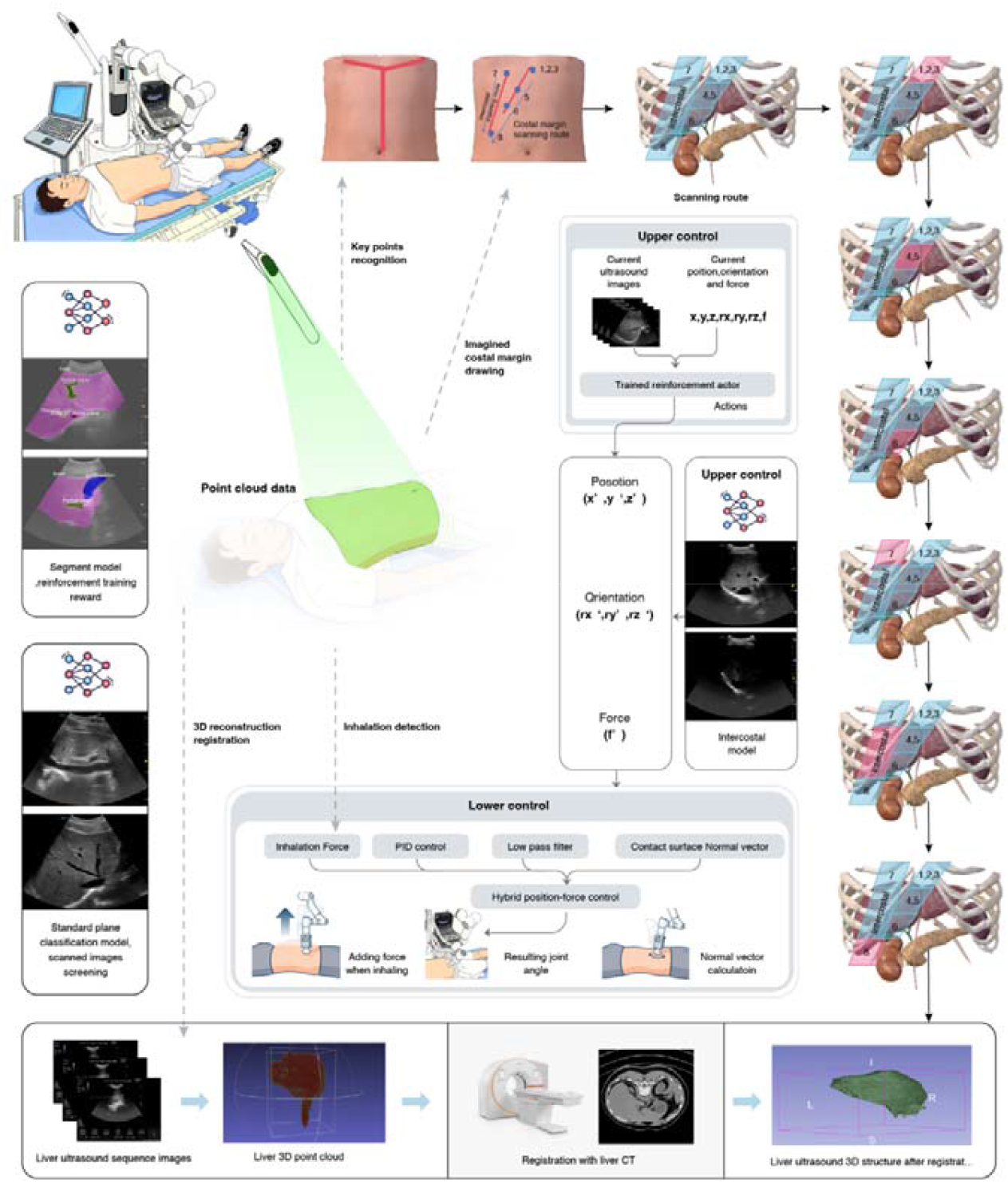
Overview of the robotic ultrasound system. Prior to scanning, the autonomous system identifies the starting point—the xiphoid process—using an RGB image recognition model. Subsequently, the 3D point cloud data of the abdominal region are acquired and utilized for dividing the scanning area and planning the baseline path. Six sub-regions are defined to encompass common liver scanning areas, including the left liver, costal margin, and intercostal regions. In each sub-region, at least one standard plane is identified. The upper-level control model transmits the subsequent target position (x, y, z) and orientation (rx, ry, rz) to the lower-level hybrid control system, thereby guiding the scanning path and ensuring appropriate contact between the ultrasound probe and the skin. Following the scanning of each sub-region, the acquired images are aligned to facilitate 3D reconstruction of the liver and computation of the overall scanning coverage.

**Figure 2.**
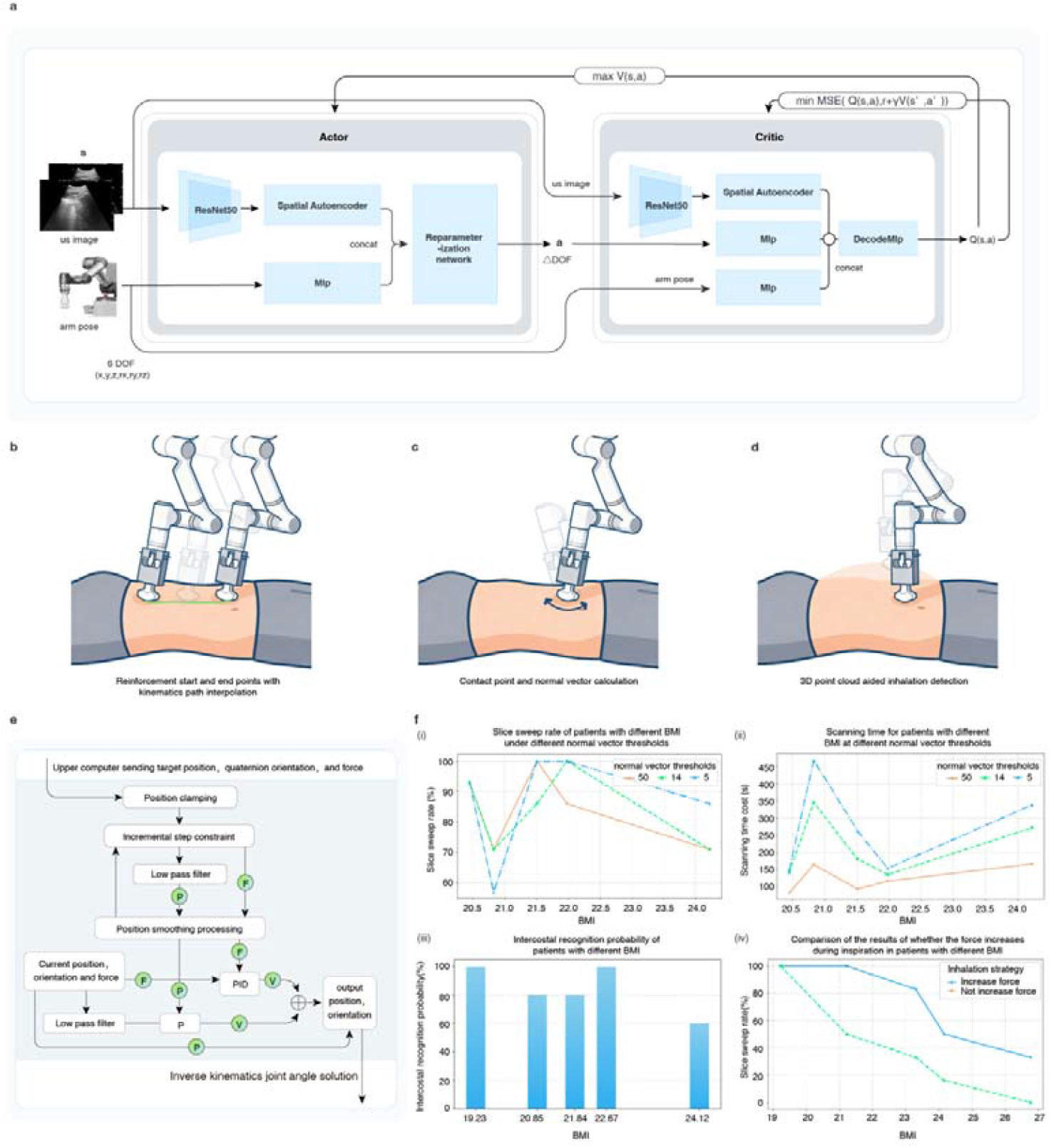
Hierarchical control architecture of the autonomous ultrasound system and ablation studies across subjects with varying body mass indices (BMIs). **a**, Reinforcement learning-based policy network for scan guidance using real-time ultrasound images. **b**, The low-pass filter ensures smooth movement towards the target points designated by the high-level policy planner without instantaneous acceleration, deceleration, reverse motion. **c**, Probe contact point and orientation adjustment using real-time normal vector estimation. **d**, Real-time respiratory (inhalation) phase detection using 3D point cloud data. **e**, Overall architecture of the lower-level hybrid force-position control system. **f(i)**, Standard plane acquisition rates across subjects with varying BMIs at different normal vector computation thresholds. **f(ii)**, Scanning durations across subjects with varying BMIs at different normal vector computation thresholds. **f(iii)**, Intercostal space recognition accuracy across subjects with varying BMIs. **f(iv)**, Comparison of standard plane acquisition rates with and without increased force application during inspiration across subjects with varying BMIs.

The ordered target positions from reinforcement actor are discrete points on human body surface (Fig. 2b). We need fluent motion control for the ultrasound robot to perform smooth and save motion records on human body surface while executing the position & orientation orders from the upper reinforcement actor. To accomplish this, we device a hybrid position and force control, which can incorporate contact point normal vector calculation (Fig. 2c) and 3D-point cloud-based inhalation detection (Fig. 2d). Ultrasound standard images are also collected on the path between reinforcement ordered points, normal vector calculation is aimed for aligning the probe’s Z-axis (probe coordinate system) with the contact surface normal vector for maintaining high image scanning quality. The inhalation detection mechanism is based on the ultrasound doctor’s habit of applying force when scanning higher BMI volunteers with excessive fat. The overall structure for the lower control framework is illustrated in Fig. 2e.

Fig. 2f(iv) shows that applying increased force while the volunteer inhales did help with the rate for collecting standard planes. The intercostal scanning procedure is less sensitive to BMI, and thus can provide stable scanning as shown in Fig. 2(iii). Applying normal vector calculation for the contact points during path interpolation achieves higher standard planes collecting rate for bigger MBI volunteers (Fig. 2(1)). However, the time cost for applying normal vector is overall high, especially for the smaller MBI volunteers. A balance can be struck by opting to activate the normal vector function when scanning volunteers with higher BMI (> 24).

#### (1) Abdominal Standard Plane Classification Model

During the inference scan, the robot performs real-time standard plane classification of liver B-mode images (9 classes in total). If the classification model does not find a standard section during the scan for a single standard plane area—20 moves of the reinforcement policy generator (a qualified standard plane is defined as the classification model’s prediction confidence level is greater than 0.8), the system will keep the top rated images for substitution. In this classification task, the metrics used are accuracy, precision, recall, and f1-score. We conducted bootstrapping for 5 times, and the results on the test set are shown in Fig. 3b(i).

**Figure 3.**
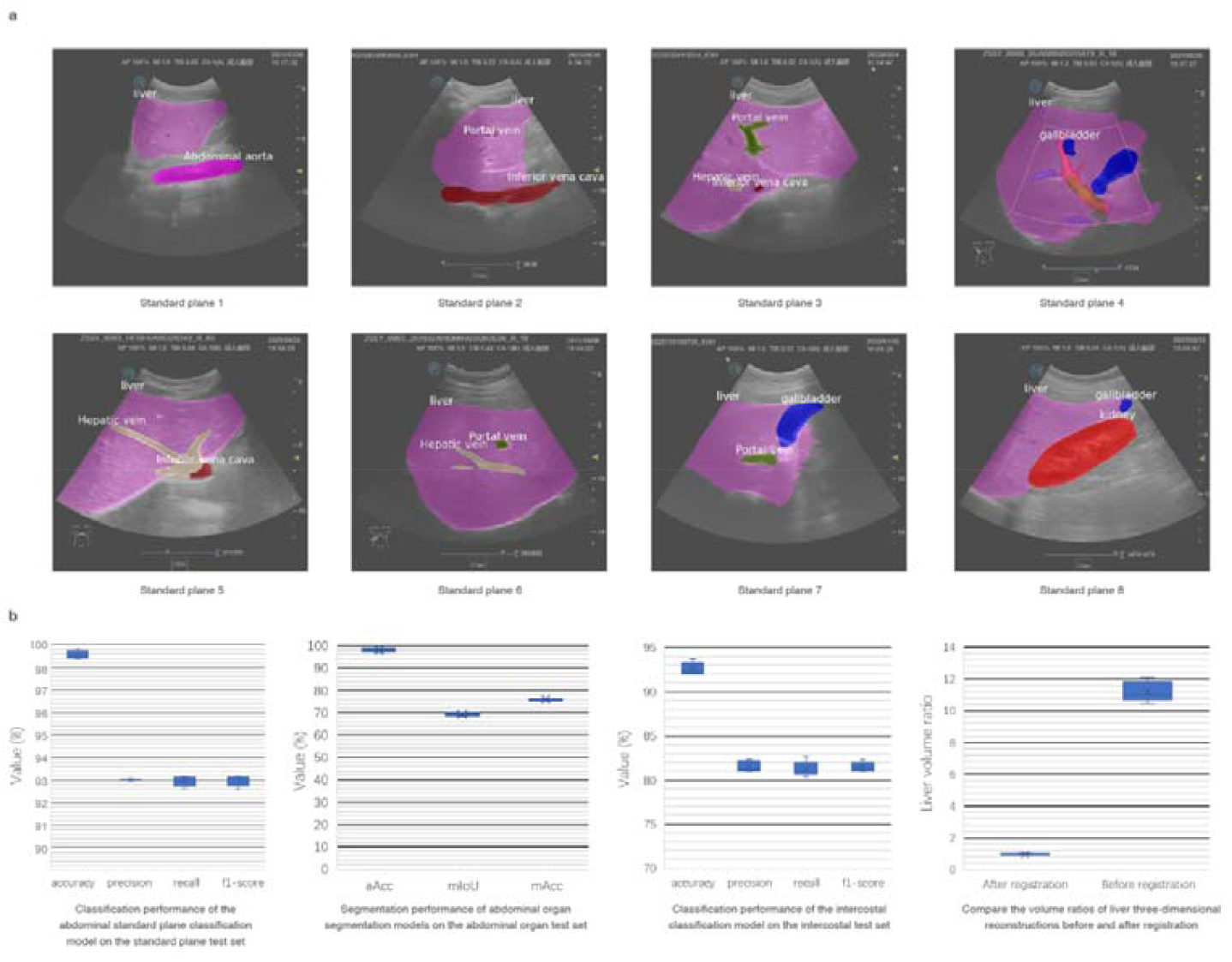
**a**, Definitions of the eight standard planes. b, Validation results for the standard plane classification model, organ segmentation model, intercostal space model, and 3D reconstruction registration model.

#### (2) Abdominal organ segmentation model

During inference scan, if the standard plane classification fails to find a standard plane, abdominal organs must be segmented. A reinforcement learning strategy guides the probe’s position and rotation angle based on the organ segmentation results. This supervised training of the organ segmentation model enables automated scanning. In the organ segmentation task, the indicators used are aAcc (average Accuracy), mIoU (mean Intersection over Union), and mAcc (mean Accuracy). This paper conducted 5 times bootstrapping, and the results on the test set are shown in Fig. 3(ii).

#### (3) Intercostal classification model

During inference scan, the intercostal areas needs to be scanned. Through supervised training of the intercostal classification model, it is possible to automatically determine whether the probe is in the optimal intercostal space based on the image. In this classification task, the indicators used are accuracy, precision, recall, and f1-score. We conducted five times bootstrapping, and the results on the test set are shown in Fig. 3(iii).

#### (4) Liver 3D reconstruction module

After complete scanning of the liver, the collected ultrasound B-mode images need to be aligned for 3D reconstruction. The process is divided into the following steps: generating point cloud data, reconstructing 3D surfaces and slicing, registering ultrasound and CT images, and visualizing the 3D reconstruction (Fig. 1(iv)). This paper uses the liver volume ratio to measure the completeness of the 3D reconstructed liver. The volumes of the ultrasound liver 3D reconstruction data before and after registration were calculated separately, while the reconstructed volume of the paired CT liver data was used as the gold standard. The liver volume ratio is defined as the reconstructed liver volume divided by the gold standard. A liver volume ratio closer to 1 indicates a closer approximation to the true liver volume. We performed five bootstrapping runs, and the results on the test set are shown in Fig. 3(iv).

## Discussion

In this study, we propose an end-to-end autonomous robotic ultrasound system designed to bridge the longstanding simulation-to-reality (sim-to-real) gap in medical robotics. Unlike prior approaches that rely heavily on phantom data or simulated environments, we train reinforcement learning (RL) policies directly using in vivo human data. This paradigm shift substantially mitigates the limited real-world generalization of conventional methods by exposing the agent to the inherent anatomical diversity and physiological complexity of living subjects during training, thereby enabling the development of highly robust scanning strategies that generalize well beyond models trained on idealized phantoms. These results validate our central hypothesis that learning from the “noise” and variability inherent in real human anatomy is a critical pathway toward clinically viable autonomous systems.

One important finding of this study is that the proposed control strategy plays a crucial role in scanning difficult subjects. Specifically, the integration of a hybrid force-position controller with real-time normal vector calibration enables the probe to maintain optimal acoustic coupling, closely mimicking the dexterous maneuvers of experienced sonographers. In addition, we introduce a respiratory phase detection mechanism that dynamically increases the applied force during inspiration, a design choice that proves decisive for imaging subjects with higher body mass index (BMI). Existing robotic ultrasound systems often struggle to acquire clear images in obese patients due to increased acoustic attenuation and thickened subcutaneous fat; in contrast, our system emulates expert behavior by applying greater pressure during inspiration to displace adipose tissue and reduce the distance to the target organ, thereby significantly improving the success rate of standard plane acquisition in high-BMI volunteers. These findings indicate that for autonomous systems to be effective in the general population, they must go beyond pure path planning and deeply integrate physiological sensing with force control strategies.

Although the standard plane acquisition rate is a commonly used metric for evaluating robotic ultrasound performance, we argue that it does not fully capture scan quality. For example, a system may reach the target plane while failing to cover diagnostically relevant regions of the organ volume. Therefore, we introduce a coverage metric based on 3D reconstruction and validate it using CT data to more comprehensively quantify system performance. Experimental results show that the proposed system achieves high liver volume coverage, demonstrating that it not only “finds” specific views but also performs more comprehensive volumetric scanning of the anatomy. This capability is critical for avoiding missed focal lesions in autonomous screening, thereby substantially enhancing diagnostic confidence.

Despite these encouraging results, several limitations of this study warrant discussion. First, although the RL policy was trained on a relatively large dataset, the volunteer cohort used for validating 3D reconstruction coverage was small (n = 13), and larger, multi-center clinical studies are needed to assess the statistical robustness and generalizability of the metric. Second, this study focuses primarily on healthy volunteers; in real clinical settings, pathological conditions such as cirrhosis or large tumors may introduce additional challenges due to altered acoustic properties and morphological deformations. Accordingly, future iterations should incorporate diverse pathological data into both training and evaluation pipelines to enhance robustness in diagnostic scenarios. Finally, while real-time normal vector computation improves contact quality, it also increases computational overhead, which may negatively affect system speed and efficiency when scanning low-BMI subjects. Developing an adaptive mechanism that activates complex control functions only when warranted by real-time image quality and task difficulty could further optimize overall scanning efficiency.

In summary, this study demonstrates that an autonomous ultrasound system combining in vivo reinforcement learning with physiologically informed hybrid control can achieve scan consistency and stability approaching expert-level performance. By effectively addressing imaging challenges arising from anatomical variability and difficult patient body types, this technology represents a substantive step toward truly operator-independent ultrasound robotics and holds promise for expanding access to high-quality diagnostic imaging in resource-limited settings.

## Methods

### Human Key Point Detection Model

For automatic scanning, it is necessary to determine the coordinates of specific human key points in advance. To this end, we trained a human key point detection model. Five anatomical key points were defined: the laryngeal prominence, left nipple, right nipple, xiphoid process, and navel. We adopted the MMPose framework [1] as the basis for our model.

A total of 3,124 RGB images of patients in the supine position were collected and manually annotated. Among them, 2,500 images were used for training, 312 for validation, and 312 for testing. The model was trained for 500 epochs with a learning rate of 0.0005 using the Adam optimizer. The mean squared error (MSE) loss was employed, and the batch size for the training set was set to 32.

### Skin segmentation model

To ensure that the scanning process was conducted exclusively on the patient’s body surface and free of obstructions, we developed a skin segmentation model to identify the accessible scanning region and facilitate safe and effective autonomous ultrasound acquisition. For this purpose, RGB images of the abdominal skin surface were collected from multiple patients in the supine position and manually annotated. A YOLOv8s-based segmentation network was employed, initialized with the pre-trained weights from yolov8s-seg.pt. Training was performed using the default hyperparameters, with the following specifications: 100 epochs, an input image size of 640 × 640 pixels, a batch size of 8, and task set to ‘segment’. A total of 500 annotated images were used, with 400 assigned to the training set, 50 to the validation set, and 50 to the test set. The trained model was subsequently applied to identify patient skin regions, thereby establishing the groundwork for safe and reliable autonomous scanning.

### Intercostal Scanning of the Left Hepatic Lobe

Intercostal scanning was performed following the completion of plane 7. The starting position for intercostal scanning was defined as the scanning start point of plane 7, shifted 8 cm along the negative direction of the y-axis. The primary objective of this step was to accurately localize an intercostal space. To this end, we trained an intercostal model capable of recognizing rib spaces from ultrasound images.

Around the defined starting position, we generated 25 candidate probe poses by applying controlled perturbations. These included five translations along the vertical axis (upward/downward), five translations along the lateral axis (left/right), five rotations about the Rz axis, and ten rotations about the Ry axis. The robotic arm sequentially navigated through these 25 poses, and at each pose the intercostal model was used to evaluate whether the acquired ultrasound image corresponded to an intercostal space. Upon positive identification, the search was terminated, and the probe orientation was further adjusted by modifying the Rx rotation angle to ±30°. The robotic arm then executed these two poses to achieve fan-shaped scanning across the identified intercostal space.

If no intercostal space was detected across all 25 candidate poses, an alternative strategy was applied. Specifically, an organ segmentation model was used to segment the liver in the ultrasound images obtained from these poses. The pose yielding the largest segmented liver area was selected as the intercostal space for subsequent fan-shaped scanning.

The entire intercostal scanning procedure was performed at two adjacent intercostal spaces. For the second intercostal space, the probe pose was directly derived from that of the first intercostal space by translating 3 cm along the negative x-axis. At this location, fan-shaped scanning was again carried out by adjusting the Rx rotation angle, thereby completing the intercostal scanning procedure.

### Force–position hybrid control

#### (1) Limiting and smoothing of the target pose and force

The target pose and force are sent by the upper computer, and the lower computer performs limiting and smoothing processing on them. For the target force, it is compared with the current force. If the current force is less than 1□N and the target force is greater than 5□N, the target force is first set to 5□N, and the smoothed target force is adjusted at a rate of 1.25□N/s. For the target position, the x-component is constrained to the range of -250 to 250□mm, and the y-component is limited to the interval of 200□mm to 500□mm. The rate of change in the target position is constrained to 50□mm/s. For the target orientation, an optimized nonlinear filter is applied for smoothing.

#### (2) Force Control

A PID controller is employed for force regulation. In the z-direction, the PID controller takes the current force and a smoothed target force as input and outputs a velocity command in the z-direction. For the x and y directions, PID control is only activated when the measured force exceeds a predefined threshold. This strategy allows the manipulator to respond to external disturbances while ensuring safety by preventing unintended motion that could harm a human operator.

Along a specified direction (determined during initialization or derived from force decomposition via sensors, rather than being commanded by the upper-level controller), the measured force is projected onto the target direction via the dot product. The resulting component, representing the force along the specified direction, is compared with the target force and processed by the PID controller to compute the velocity along that direction. This velocity is then decomposed into its x, y, and z components.

The system employs a hierarchical PID control strategy to maintain constant contact force between the ultrasonic probe and the tissue while adjusting its orientation. Compared to admittance control, the PID controller generates control signals directly from the proportional, integral, and derivative terms of the error, offering advantages such as structural simplicity, clearly defined parameter meanings, and high computational efficiency. The control framework consists of a force control loop, which maintains a constant contact force, and an orientation control loop, which adjusts the probe’s alignment. Joint commands for the robot are generated through inverse kinematics mapping.

Among them, the objective of the force control loop is to maintain constant force contact *f*_d_ between the probe and the tissue. The error *e*_f_ between the current contact force *f*_z_ and the target value *f*_d_ is input into a PID controller, which calculates the correction amount for the Z-axis position.

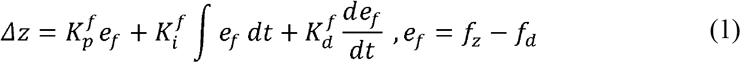

The parameters 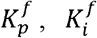, and 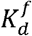 represent the proportional, integral, and derivative gains, respectively, which were tuned experimentally to optimize the dynamic response. In the practical implementation, the integral term accumulation was frozen when the actuator reached its saturation limit to prevent overshoot, while a 20 Hz low-pass filter was applied to the derivative term to suppress high-frequency noise.

#### (3) Position Control

### A proportional (P) controller is employed for position regulation

To mitigate noise, the current position is filtered prior to PID control to reduce interference. The difference between the smoothed target position and the filtered current position is fed into the P controller to compute the output velocity, with the proportional gain set to P=1. Finally, the velocity outputs from the force control and position control loops are summed to yield the final commanded velocity. This final velocity is used to update the target position, which, together with the target orientation Q, is then input to an inverse kinematics solver to compute the corresponding joint angles for the robotic arm.

### Contact Point Normal Vector Estimation

To enable adaptive probe orientation adjustment and safe contact control during ultrasound examinations, while ensuring both imaging quality and patient comfort, the system employs a contact force sensing algorithm based on segmented plane modeling and torque-based inverse computation. Using six-axis force/torque sensor data, the algorithm estimates the contact point position and the normal direction between the robot end-effector and the tissue surface. These estimates are then fed back to the control module to achieve dynamic posture adjustment.

During ultrasound imaging procedures, the interaction between the probe and tissue involves a complex, non-uniform pressure distribution. To simplify modeling, the equivalent mechanics principle is applied, whereby the actual distributed forces are approximated as a resultant force and torque acting at a single contact center point, as illustrated in Fig. 2c the model realizes force perception through the following steps:

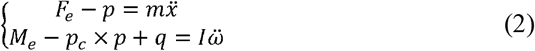

Here, *m* represents the probe mass, and *I* denotes the moment of inertia. *q* is the local moment generated by deformation of the contact surface, while 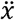 and 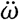 represent linear acceleration and angular acceleration, respectively. When the ultrasound probe acquires images, the system is in a quasi-static state ( 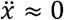 and 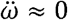). Under these conditions, the equation can be simplified as:

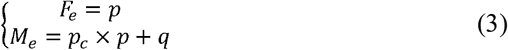

In contact force modeling, the direction of the contact moment is determined by establishing the parallelism (*q = k∇S*(*p*_*c*_)) between the local moment and the normal vector of the contact surface, and constructing a set of piece-wise equations ({*S*_1_, *S*_2_, …, *S*_*n*_}) representing the instrument surface through a multi-plane fitting method.

To represent the irregular surface between the probe and the contact object, a predefined set of triangular patches is employed to discretely model the contact region. Each patch is defined by three points *p*_1_ *p*_2_ *p*_3_ ∈ ℝ^3^, its corresponding plane equation is constructed by calculating the normal vector via the cross product of vectors *n =* (*p*_3_ − *p*_1_) × (*p*_1_ − *p*_1_).

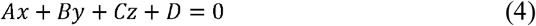

Where [A, B, C]^⊤^.= n, D =m −n^⊤^ p_1_. Each triangular facet is associated with a corresponding normal vector and plane parameters. Upon receiving the latest six-dimensional force/moment data f = [F_x_, F_y_, F_z_, M_x_, M_y_, M_z_]^T^ the system iterates through all triangular facets sequentially, constructing the following 4×4 moment equilibrium constraint equation system for each facet.

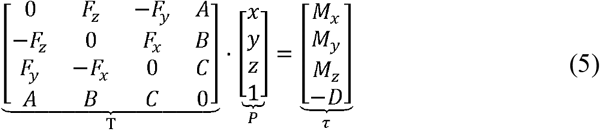

Solving the system of equations yields the theoretical contact point *P* = [*x, y, z*]^⊤^on the current surface patch. The barycentric coordinate (*u, v, w*) method is then employed to determine whether the point lies inside the triangle. This is achieved by computing the barycentric coefficients via vector dot products. If (*u, v, w*)≥0 and *u* + *v* + *w* = 1 are satisfied, the point is considered to lie within the surface patch. For all candidate solutions located inside the patch, the cosine of the angle 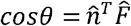 between the normal vector 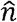 and the unit force direction 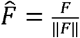 is further calculated. The solution corresponding to the smallest angle—i.e., the one most aligned with the force direction—is selected as the final estimated contact point.

### Ballooning Detection and Dynamic Force Adjustment

To achieve precise and stable ultrasound scanning, the system first performs surface modeling of the target contact region using point cloud mapping. Specifically, a structured-light depth camera is employed to acquire three-dimensional point cloud data of the probe’s scanning area, yielding discrete points with spatial coordinates (*x, y, z*). After Voxel grid filtering, the point cloud is projected onto the Y–X plane to generate a two-dimensional height map, where each grid cell records the mean height z_map_ of the local point cloud as its representative elevation value. During operation, the system continuously acquires the end-effector’s current pose along with the pressure compensation signal, and uses these inputs to construct the following ballooning detection formula:

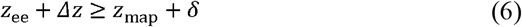

A fixed threshold*δ*, empirically determined (set *δ* = 5/8*mm* in this system), was applied. Once the condition in Equation (9) was satisfied, the system identified the occurrence of abdominal expansion and dynamically adjusted the desired contact force accordingly.

### Reinforcement-learning framework

We adopted the refined Soft Actor-Critic (SAC) algorithm introduced in Ref. 1 as our core reinforcement-learning (RL) architecture. This off-policy, maximum-entropy deep-RL method retains the stability and exploration benefits of entropy maximization while simultaneously improving sample efficiency—an essential requirement when the patient morphology space is large and heterogeneous, as in free-hand ultrasound.

### RL environment and policy network

The RL environment is formally defined by a three-element tuple: (i) a real-time ultrasound frame acquired from the clinical scanner, (ii) the seven-dimensional pose vector of the robotic arm, and (iii) the canonical-plane class label (scalar). To accommodate this multimodal state space, we designed a bespoke policy network. A pretrained ResNet-50 extracts spatial features from the ultrasound image; a shallow multi-layer perceptron (MLP) encodes the arm pose; and a learnable embedding maps the target plane label into a latent vector. The three representations are concatenated and fed to a Tanh-Gaussian stochastic policy head that outputs the parameters of a seven-dimensional Gaussian distribution. An action sampled from this distribution specifies the incremental pose adjustment applied to the robot arm.

### Value network architecture

The value function approximator mirrors the encoder structure of the policy network. Ultrasound frames are embedded via ResNeXt-50; the seven-dimensional pose is processed by an MLP; the chosen action is similarly encoded by a separate MLP; and the target plane label is again embedded. The four resulting vectors are concatenated and decoded by a final MLP that regresses the state-action value.

### Loss terms

We adopt the typical SAC loss terms that comprises of the following three objectives:

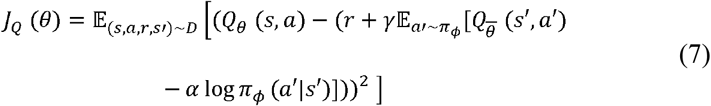

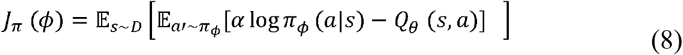

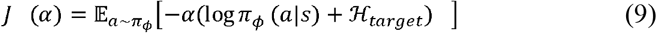

Where (*s, a*) and (*s′, a′*) are the current and next state and action pairs, θ and ϕ is the Q function and the policy network parameters. α is the temperature parameter that balances the entropy and the rewards. ℋ _*target*_ is the target entropy and is set to the negative value of the action space dimension.

#### (1) Reward Function

The primary objective of this scanning task is to localize eight standard ultrasound (US) views of the liver. To enhance accuracy, we extracted anatomical features from US images of each standard view. An organ segmentation model was applied to delineate anatomical structures within each view. For each standard view, 100 US images were selected to compute the average area of segmented organs. Based on these statistics, weights were assigned to each organ using an inverse relationship and subsequently normalized to obtain the relative area proportions for each standard view.

The reward function was then defined based on the classification probability and segmentation results of consecutive US frames. Specifically, let prob denote the predicted probability of the current view classification, *a*_*curr*_ the segmented organ areas in the current frame, *a*_*last*_ those in the previous frame, and weight the normalized organ area proportion for the corresponding standard view. The reward was computed as follows:

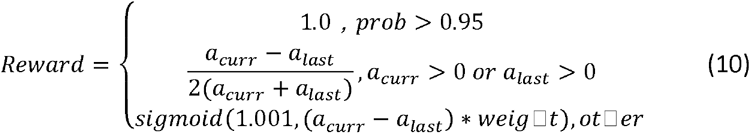

where *sigmoid* (□_1_,□_2_) is defined as

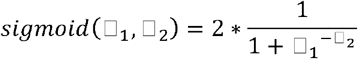

#### (2) Training Procedure

We trained separate models for each of the eight standard liver views. The following hyperparameters were used: policy learning rate = 1e-4, soft update coefficient = 5e-3, Q-function learning rate = 1e-4, and maximum path length = 11. Training was conducted on a large cohort of volunteers, resulting in eight distinct weight sets one for each standard view which were subsequently used for inference during the scanning process.

#### (3) RL-Based Scanning Inference Pipeline

Upon completion of training, the inference pipeline proceeds as follows. First, the trained RL model weights are loaded, and the robotic arm is moved to the initial scanning position for the target view. The current US image and robotic pose, along with the target view class, are input to the trained policy network. The network outputs an action, which is added to the current pose to compute a new target pose. This target pose is sent to the robotic arm, which moves accordingly. The process is repeated iteratively, enabling the RL agent to guide the robotic arm through the scanning trajectory.

Each view scan proceeds for a maximum of 20 steps. After every 10 steps, the robotic arm is reset to the initial scanning position for the current view. The entire scanning session consists of sequentially scanning all eight standard liver views. A view is considered complete either when the maximum number of steps is reached or when the view classifier outputs a confidence score above a predefined threshold.

### Standard plane Classification Model

#### (1) Data acquisition and preprocessing

In this study, a total of 16,866 B-mode abdominal images were collected using a Clover portable ultrasound device, and the standard planes were annotated by two ultrasound physicians. The number of annotated standard planes is shown in Table 2. We also collected images of non-standard planes and named them standard plane 0. Image preprocessing included subtracting the mean and dividing by the standard deviation of the three channels. The means of the three channels were 122, 116, and 104, and the standard deviations were 68, 66, and 70.

**Table 2.**
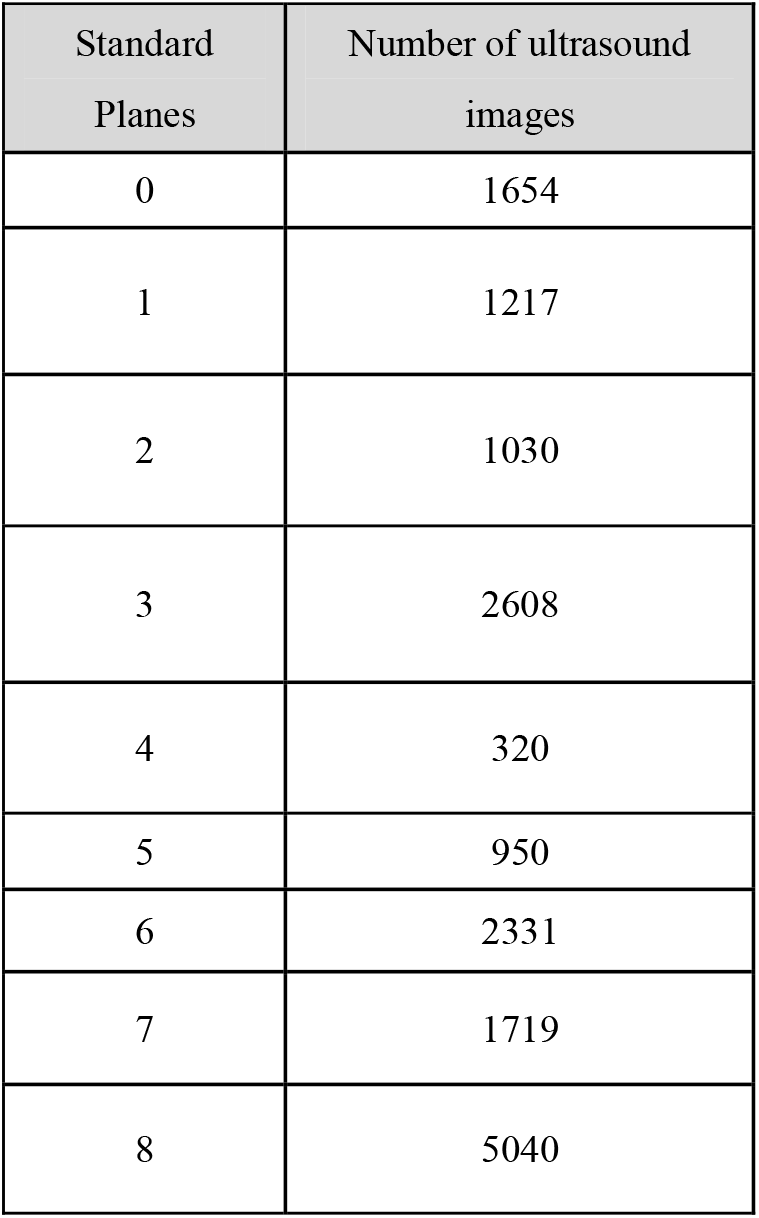
The categories of standard planes and the number of ultrasound images used for training in the liver standard plane classification model.

| Standard Planes | Number of ultrasound images |
| --- | --- |
| 0 | 1654 |
| 1 | 1217 |
| 2 | 1030 |
| 3 | 2608 |
| 4 | 320 |
| 5 | 950 |
| 6 | 2331 |
| 7 | 1719 |
| 8 | 5040 |

#### (2) Training process

The classification model structure used in this paper is the ResNeXt101-32x8d model [20]. In the backbone part, the depth is set to 101, the group is set to 32, and each group is assigned 8 channels. At the same time, the pre-trained weights (open-mmlab://detectron2/resnext101_32x8d) are used. In the head part, the number of input channels is set to 2048, and the number of output channels is set to 9. The dataset is divided into 15181 (train set), 1685 (valid set), and 950 (test set). The epoch is 100, the learning rate is set to 0.001, the optimizer is SGD, the loss function is Focal loss [21], and the batch size of the training set is set to 40.

### Abdominal organ segmentation model

#### (1) Data acquisition and preprocessing

In this paper, a total of 12,190 B-mode abdominal images were collected using a Clover portable ultrasound device, and two ultrasound physicians performed multi-organ contour annotation. The organ categories and corresponding numbers are shown in Table 3. Image preprocessing includes subtracting the mean and dividing by the standard deviation of the three channels, and scaling the image to 512x512 size.

**Table 3.** The organ type and the number of ultrasound images used to train the segmentation model.

| Organ Category | Number of organs |
| --- | --- |
| Liver | 3977 |
| Portal vein | 1616 |
| Gallbladder | 1360 |
| inferior vena cava | 1201 |
| Hepatic vein | 1190 |
| Kidney | 918 |
| bladder | 621 |
| Pancreas | 608 |
| Spleen | 540 |
| Abdominal aorta | 377 |
| Splenic vein | 260 |
| Other blood vessels | 49 |
| Common bile duct | 4 |
| Posterior rib shadow | 5739 |
| Transverse cut of ribs | 4465 |
| Longitudinal section of the ribs | 1471 |

#### (2) Training process

The segmentation model structure used in this paper is the DeeplabV3 plus model [22]. The depth of the backbone is set to 50, the number of input channels of the decoder head is set to 2048, and the number of output channels is 17. At the same time, pre-trained weights (open-mmlab://resnet50_v1c) are used. The input channels of the auxiliary head are set to 1024, and the number of output channels is set to 17. The dataset is divided into 10791 (train set), 1399 (valid set), and 579 (test set). The epoch is 100, the learning rate is set to 0.01, the optimizer is SGD, the loss function is Cross Entropy loss, and the batch size of the training set is set to 20.

### Intercostal classification model

#### (1) Data acquisition and preprocessing

A total of 6980 B-mode abdominal rib intercostal images were collected using a Clover portable ultrasound device and annotated by two ultrasound physicians (intercostal or not intercostal). The number of annotations is shown in Table 4.

**Table 4.** The intercostal and non-intercostal images are used to train the recognition model.

| Category | Number |
| --- | --- |
| Intercostal | 3411 |
| Not in the intercostal space | 3569 |

#### (2) Training process

This article uses the ResNeXt50 classification model. In the backbone, the depth is set to 50, the group is set to 32, and each group is assigned 8 channels. Pretrained weights (open-mmlab://detectron2/resnext50_32x8d) are also used. In the head, the number of input channels is set to 2048, and the number of output channels is set to 2. The dataset is split into 6282 (train set), 698 (valid set), and 3077 (test set). The epoch number is 15, the learning rate is set to 0.001, the optimizer is SGD, the loss function is Cross Entropy loss, and the batch size of the training set is set to 128.

### Liver 3D reconstruction module

#### (1) Data acquisition and preprocessing

We recruited 13 male volunteers, each of whom used the robot to collect about 10,000 B-mode abdominal ultrasound images, as well as the liver region segmentation mask and probe posture information (position and posture) recorded by the robotic arm for each ultrasound image. CT liver data was also collected from these volunteers.

#### (2) Generate point cloud data

First, we need to extract the pixel points on the edge of the liver segmentation mask and obtain the coordinates (u, v) on the image plane. Then, for each edge pixel point, we calculate the position relative to the ultrasound origin on the image plane based on the imaging parameters on the ultrasound image, see equation (11).

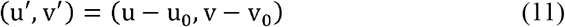

(u_0_, v_0_) is the position of the ultrasound origin in the image coordinate system, and u′,v′ is the coordinate of the edge point in the ultrasound origin coordinate system. Then, according to the depth parameter of the ultrasound device (the actual physical length δ corresponding to each pixel), the pixel coordinates are converted into the actual physical coordinates in the probe coordinate system, see equation (12).

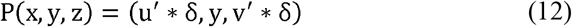

Where P (x,y,z) represents the physical coordinate system under the probe coordinate system, and x, y, and z represent the values on the x-axis, y-axis, and z-axis. Since ultrasound is a cross-sectional detection, the value on the y-axis is always 0.

Finally, the probe pose T_probe_ provided by the robotic arm is used to transform the liver edge point from the probe coordinate system to the robotic arm coordinate system (i.e., the world coordinate system), see equation (13).

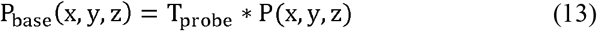

P_base_ (x,y,z) represents the position of the liver edge point corresponding to each ultrasound frame in the robotic arm coordinate system. For all frames in the ultrasound sequence, the corresponding P_base_ (x,y,z) can be mapped to the same coordinate system to obtain the complete liver point cloud data.

#### (3) 3D surface reconstruction and slice processing

After obtaining the complete liver point cloud data, it is necessary to voxelize the point cloud, then use the Marching Cubes algorithm to generate a triangular mesh, smooth and fill the holes in the mesh, and finally reconstruct the 3D surface. Along the z-axis direction, slice the voxelized liver point cloud with a spacing of 0.001 voxel size between each slice, and each slice represents a cross-section of the liver point cloud along the z-axis

Registration and 3D reconstruction visualization between ultrasound and CT images: There are some discrepancies between the liver point cloud data generated based on ultrasound images and the structure of the real liver, while the liver structure scanned by CT is more complete. Therefore, a registration algorithm is needed to map the ultrasound liver point cloud slice data to the CT liver data, so as to achieve a closer match to the real liver structure. The registration algorithm used in this paper is DeepReg [23]. During the training phase, the input data is the paired ultrasound liver point cloud slice data (as a moving image) and CT liver data (as a fixed image) of the same volunteer. The registration network will output a dense displacement field (DDF) with the same shape as the moving image. Each value in the DDF can be regarded as the position of the corresponding pixel in the moving image. Therefore, the DDF defines the mapping from the moving image coordinates to the fixed image. During the training process, the loss used is Dice loss and Bending loss, the optimizer is Adam, the learning rate is set to 1.0×10^−5^, the batch size is 4, and the epoch is 5000. The training set is the paired data of 8 volunteers, the validation set is 3, and the test set is 2.

During the inference phase, the input data is ultrasound liver point cloud slice data and the trained DDF weights. The output is the registered ultrasound liver point cloud slice data. The slice data is then binarized and VTK (visualization toolkit) is used to reconstruct the 2D slice data into 3D to obtain a 3D liver visualization structure. The spacing in the x and y directions is set to 1 pixel, and the spacing in the z direction is set to 2.5 pixels.

## Acknowledgements

All experiments with human participants were performed with the approval of The Institutional Review Board of BGI (BGI-IRB 24080). The authors are grateful to the participants and explained the full process to them, and each signed an informed consent form (including their understanding of publishing information in the journal). The authors wish to thank medical personnel from the Department of Ultrasound of Shanghai Zhongshan Hospital for their invaluable cooperation.

## Author Contributions

The manuscript was prepared with contributions from all authors. All authors have approved the final version of the manuscript.

## Notes

### Competing Interest Statement

The authors have declared no competing interest.

